# Multiscale superinformation reveals structured organization of genomic sequence features

**DOI:** 10.64898/2026.11.22.768374

**Authors:** Sarvesh Galgale, Praharshit Sharma

## Abstract

Genomic sequence organization varies across spatial scales, but sequence-complexity measures are often evaluated at a single resolution. Superinformation (SI), introduced by Bose and Chouhan, measures heterogeneity in local sequence entropy by taking the entropy of the distribution of block-level Shannon entropy values. We characterize SI across 14 Sequence Ontology-defined genomic feature classes in seven species and seven block sizes from 20 to 200 bp. SI profiles were non-increasing for 72 of 86 evaluable species-feature combinations, although this pattern was not universal. Observed SI generally exceeded both a block-count-controlled random null and a species-level GC-controlled null. Associations with mononucleotide entropy, dinucleotide entropy, and GC content were modest, with no individual descriptor explaining more than 10.4% of the observed variation. For the 13 complete feature-level profiles used in geometric analysis, the first two principal components explained 97.859% of the variance, and hierarchical clustering grouped features with similar SI trajectories. We also summarize the profiles using the Characteristic Information Scale and Information Persistence Score. These analyses extend the original super-information framework to a comparative, multiscale analysis of genomic feature heterogeneity while keeping the interpretation of SI limited to the sequence properties measured here.

## 1 Introduction

Genomic DNA is organized across multiple spatial scales. Nucleotide composition, local sequence patterns, repetitive elements, regulatory signals, and other sources of heterogeneity can occur within the same genomic region. A statistic measured at one resolution therefore gives only one view of that organization. Information-theoretic and sequence-complexity methods have been used at several scales, from global nucleotide and *k*-mer composition to local entropy, compositional domains, and sequence-context measures. Local entropy and compositional analyses show that sequence properties can vary along genomes, and comparative studies show that the useful resolution of a sequence representation can also depend on scale (Bussi et al., 2021; Cristadoro et al., 2018; Moya et al., 2020; Thanos et al., 2018).

Shannon entropy is a natural starting point for describing variability in a symbolic sequence. For DNA, nucleotide-frequency entropy summarizes uncertainty in the four-letter alphabet and is widely used to describe sequence composition and local variability (Shannon, 1948). A single entropy value, however, does not show how local information content is distributed within a sequence. Two regions can have similar nucleotide frequencies and similar entropy values while differing in how variable and compositionally constrained segments are arranged. Methods based on local entropy retain some of this spatial information, while entropic segmentation identifies domains with different local sequence properties (Moya et al., 2020; Thanos et al., 2018). Sequence comparisons can also change with representation scale, including the length of the *k*-mers used to characterize genomes (Bussi et al., 2021).

Within this information-theoretic framework, Bose and Chouhan (2011) introduced superinformation as a measure of the heterogeneity of local sequence entropy. Their construction begins by partitioning a DNA sequence into blocks of length *B*, calculating the Shannon entropy of each block, and then calculating the entropy of the resulting distribution of block-level entropy values. In this formulation, superinformation can be written as

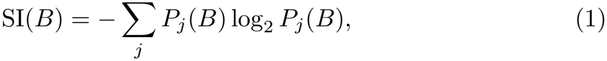

where *P*_*j*_(*B*) is the fraction of blocks whose Shannon entropy falls within entropy bin *j*. Bose and Chouhan described this quantity as an “entropy of entropy” or “randomness of randomness”, emphasizing that it measures variation among local information states rather than average nucleotide variability alone. They also treated the block size *B* as the resolution at which sequence organization is examined. Their original analysis demonstrated that superinformation could distinguish coding and noncoding regions of human DNA and could also be applied to untranslated regions, establishing its potential as a descriptor of sequence heterogeneity beyond conventional sequence averages (Bose & Chouhan, 2011).

The original framework establishes the measure, but its use across a broad set of genomic feature classes has not been examined systematically. The multi-scale literature also makes clear that local entropy, compositional segmentation, and *k*-mer analyses describe different aspects of sequence organization and can depend on the scale of analysis (Bussi et al., 2021; Cristadoro et al., 2018; Moya et al., 2020; Thanos et al., 2018). This leads to a broader question: do genomic feature classes show reproducible SI profiles across resolutions and species, and can their differences be distinguished from simple sequence-level properties? Standardized feature definitions make this comparison possible across different genomic annotations. The Sequence Ontology provides such a vocabulary for describing genomic sequence features (Eilbeck et al., 2005).

Here, we characterize superinformation across 14 Sequence Ontology-defined genomic feature classes in seven species and seven block sizes from 20 to 200 bp. We compare SI magnitude and its change with resolution, then test the observed values against complementary random null models and against mononucleotide entropy, dinucleotide entropy, and GC content. We also examine the resulting SI profiles in low-dimensional and hierarchical feature spaces and summarize their scale-dependent behavior with the Characteristic Information Scale and Information Persistence Score. The goal is not to redefine superinformation, but to examine what the existing measure captures when applied systematically across genomic features, species, and resolutions.

## 2 Methods

### 2.1 Genomic datasets and feature extraction

We analyzed reference genomic sequences from seven model organisms: human, mouse, rat, zebrafish, fly, worm, and yeast. Genome assemblies and corresponding GFF3 annotations were obtained from Ensembl release 111 (January 2024; Yates et al., 2026). The assemblies used were GRCh38 for human, GRCm39 for mouse, mRatBN7.2 for rat, GRCz11 for zebrafish, BDGP6.46 for fly, WBcel235 for worm, and R64-1-1 for yeast. Sequence Ontology annotations were used to define the genomic feature classes analyzed in this study.

Fourteen feature types were retained for analysis: CDS, exon, five-prime UTR, three-prime UTR, lncRNA, ncRNA gene, miRNA, snRNA, snoRNA, rRNA, tRNA, mRNA, pseudogene, and transposable element. Feature-specific sequences were extracted from the annotated genomic coordinates for each species.

Sequence records that could not contribute to the required block sizes were excluded. Because feature availability and sequence length differed among genomes, not every feature was represented in every species or at every block size. The resulting Superinformation dataset contained 578 feature-species-block-size observations spanning the seven species, 14 Sequence Ontology feature classes, and seven block sizes used in the study. The final feature and sequence quality-control summaries are retained with the project outputs.

### 2.2 Block-based sequence representation

For each sequence and block size *B*, the sequence was partitioned into non-overlapping blocks of length *B*. The number of complete blocks was calculated as ⌊*L/B*⌋, where *L* is the sequence length. Only these complete blocks were included in the entropy calculation; any terminal sequence remainder shorter than *B* was excluded. Blocks were analyzed independently. We examined block sizes of 20, 40, 60, 80, 100, 150, and 200 bp.

This range was selected to characterize how sequence heterogeneity changes as block size increases and the representation becomes progressively coarser. The use of block size as an explicit resolution parameter follows the original superinformation framework of Bose and Chouhan.

### 2.3 Shannon entropy of individual blocks

Sequence records containing ambiguous or non-canonical nucleotide symbols were excluded before block construction. Remaining records were partitioned into non-overlapping complete blocks of the specified sizes. The frequencies of the four nucleotides within each block were then calculated. Shannon entropy was computed as:

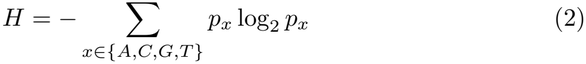

where *p*_*x*_ is the frequency of the nucleotide *x* within the block.

### 2.4 Superinformation

Superinformation was calculated following the entropy-of-entropy construction introduced by Bose and Chouhan. For a sequence divided into blocks of size *B*, let *H*_*i*_(*B*) denote the Shannon entropy of block *i*. The distribution of these block-level entropy values was then represented using a fixed 50-bin histogram spanning the theoretical nucleotide-entropy range of 0 to 2 bits.

If *P*_*j*_(*B*) denotes the fraction of blocks whose entropy falls in histogram bin *j*, Superinformation was calculated as:

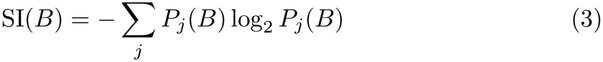

Thus, Shannon entropy measures nucleotide variability within individual blocks, whereas Superinformation measures how that local variability itself is distributed across blocks. This is the quantity referred to by Bose and Chouhan as an entropy of entropy.

The same calculation was applied at each block size, producing a multiscale SI profile for each feature-species combination.

### 2.5 Multiscale SI analysis

Superinformation profiles were evaluated across the seven block sizes of 20, 40, 60, 80, 100, 150, and 200 bp. For each species-feature combination, the sequence was therefore represented by a multiscale SI profile:

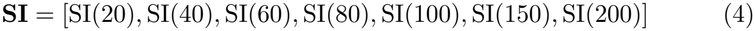

To compare the shape of profiles independently of their absolute SI magnitude, each profile was additionally normalized to its value at 20 bp:

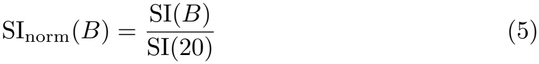

This normalization preserves the relative change in SI across the investigated block-size range while setting the 20-bp value to 1.

Initial SI decay was quantified for each species-feature combination using the slope between the first two evaluated block sizes, 20 and 40 bp:

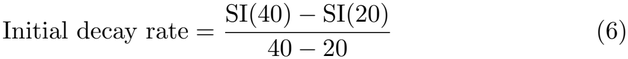

The resulting quantity has units of SI bits per bp. Negative values therefore indicate a decrease in SI between 20 and 40 bp, with more negative values representing a steeper initial decline. Combinations with fewer than two available block sizes were excluded from this calculation.

Because the number of available blocks decreases as block size increases, SI estimates at larger block sizes may be based on fewer observations. All multiscale comparisons were therefore performed using the same predefined block-size grid and the same SI estimation procedure.

### 2.6 Null models

We used complementary random null models to test whether the observed SI profiles could be reproduced under simplified sequence-generation assumptions.

#### Block-count-controlled random null

For each species, feature, and block size, independent random DNA blocks were generated in a number equal to the observed number of blocks contributing to the corresponding SI estimate. Each block was generated by sampling A, C, G, and T with equal probability. Feature-species-block-size combinations with fewer than 50 observed blocks were excluded from this null analysis. Superinformation was then calculated using the same 50-bin procedure as for the observed data.

#### GC-controlled null

For each species and block size, 50,000 independent random DNA blocks were generated using the species-level GC fraction to define the nucleotide probabilities, with A and T assigned equal probability and C and G assigned equal probability. Superinformation was calculated from the resulting entropy distribution.

#### Fast null

A computationally efficient Monte Carlo approximation was generated independently for each block size using 50,000 random blocks with uniform nucleotide probabilities. The resulting SI estimates were compared with the block-count-controlled random null to assess agreement between the two computational approaches.

For all null comparisons, the primary quantity of interest was the difference between observed and null SI at the same resolution. No explicit random seed was set for the Monte Carlo null simulations; the saved null-model tables used for the reported analyses are retained as final analysis outputs.

### 2.7 Baseline sequence statistics

To determine whether feature-level SI variation could be reduced to conventional sequence statistics, we compared mean SI with three baseline descriptors: mononucleotide Shannon entropy, dinucleotide entropy (*k* = 2), and GC content. For each Sequence Ontology feature class, mean SI was calculated across the available observations in the 578-observation SI dataset. Baseline sequence descriptors were calculated separately using 89 species–feature combinations, with up to 1,000 sequences included per combination. Associations between the resulting feature-level summaries were quantified using Pearson correlation, and the coefficient of determination (*R*^2^) was calculated as the square of the correlation coefficient.

### 2.8 Feature-space construction and dimensionality reduction

For geometric analyses, superinformation values were averaged across the available species for each Sequence Ontology feature at each block size, producing one multiscale profile per feature. The study contained 14 feature types overall; 13 had complete values at all seven evaluated block sizes and were retained for geometry, while the incomplete tRNA profile was excluded. The resulting feature-by-block-size matrix therefore had dimensions 13× 7. Missing values were not imputed or interpolated.

Principal component analysis was performed directly on the retained SI values without prior scaling so that differences in SI magnitude across resolutions were preserved. PCA was implemented using `sklearn.decomposition.PCA` with two components.

UMAP was used as a complementary nonlinear visualization of the same 13 complete feature-level SI profiles. The final two-dimensional analysis used `n neighbors=10`, `min dist=0.2`, Euclidean distance, and `random state=42`. The coordinates used in the final figure were regenerated as the canonical visualization from the frozen, unscaled 13 ×7 feature-level SI matrix and were used descriptively.

### 2.9 Hierarchical clustering

For hierarchical analysis, the same unscaled 13 ×7 feature-level SI matrix was used. SI values had been averaged across available species for each feature at each block size, and the incomplete tRNA profile had been excluded.

The resulting feature profiles were clustered using average-linkage hierarchical clustering with Euclidean distance. Four clusters were identified using the `maxclust` criterion. The saved feature assignments were used directly for the cluster-membership visualization; cluster identity was not treated as a biological or ontology classification.

### 2.10 Characteristic Information Scale

We defined the Characteristic Information Scale (CIS) operationally as the block size at which superinformation reaches 50% of its value at 20 bp. Thus, for an SI profile SI(*B*),

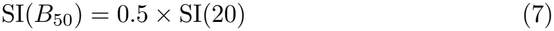

where *B*_50_ denotes the 50% threshold crossing. We refer to this quantity as CIS.

When the 50% threshold fell between two sampled block sizes, *B*_50_ was estimated by linear interpolation between the adjacent block sizes. When the threshold was not reached within the investigated range of 20 to 200 bp, CIS was recorded as not reached within the study window. This definition is based on the observed SI trajectory and does not imply a biological characteristic scale beyond the resolution range examined.

CIS was calculated independently for each available species-feature trajectory. For each feature class, finite CIS values were then summarized across available species as mean ± standard deviation (SD). Singleton estimates are reported without a standard deviation.

### 2.11 Information Persistence Score

The Information Persistence Score (IPS) was calculated as the area under the superinformation-versus-block-size curve over the available evaluated block sizes within the 20–200 bp study window:

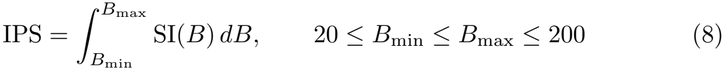

Numerical integration was performed using the trapezoidal rule across the available ordered values drawn from 20, 40, 60, 80, 100, 150, and 200 bp:

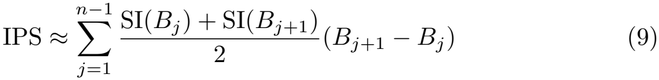

A normalized IPS was additionally calculated after scaling each SI profile by its value at 20 bp:

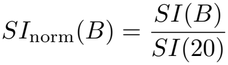

followed by the same available-point trapezoidal integration. Complete trajectories span 20–200 bp. Incomplete trajectories were integrated only over their available evaluated block sizes within that window and were not interpolated or extended to 200 bp. Because SI is expressed in bits and block size in base pairs, raw IPS has units of bit·bp. The normalized SI profile is unitless, so normalized IPS has units of bp. Raw IPS therefore reflects both SI magnitude and persistence over the available evaluated span, whereas normalized IPS emphasizes the shape and persistence of the profile after normalization to the 20-bp value. IPS was calculated independently for each available species-feature trajectory. For each feature class, raw and normalized IPS values were then summarized across available species as mean ± standard deviation (SD).

### 1.12 Statistical analysis and reproducibility

All analyses were performed in Python using NumPy, pandas, SciPy, scikit-learn, UMAP, Biopython, and Matplotlib. Statistical summaries and publication figures were generated from the final analysis outputs.

The reusable computational implementation was separately packaged and validated against retained reference outputs. The packaged CIS implementation reproduced the reference CIS dataset with identical row structure, identical finite-value coverage, and no NaN-pattern mismatches. Among 22 finite CIS values compared between the reference and reproduced datasets, the maximum absolute difference was 1.42 × 10^*−*14^.

## 3 Results

### 3.1 Superinformation reveals feature-dependent multiscale sequence heterogeneity

We calculated superinformation (SI) for 14 Sequence Ontology feature classes across seven species and seven block sizes from 20 to 200 bp. At each resolution, SI was obtained from the distribution of Shannon entropy values across sequence blocks. This applies the block-based entropy-of-entropy measure introduced by Bose and Chouhan to a broader comparison of genomic feature classes.

SI magnitude differed among feature classes. At 20 bp, three-prime UTR, lncRNA, and five-prime UTR had among the highest mean values, at 3.82, 3.80, and 3.73, respectively. CDS had a lower mean of 3.31, while tRNA was among the lowest at 2.96.

Across the complete profiles, the same separation remained. Averaged across available species and block sizes, lncRNA had the highest mean SI at 2.96, followed by three-prime UTR at 2.95, five-prime UTR at 2.87, ncRNA gene at 2.87, and mRNA at 2.83. Lower mean values occurred for CDS at 2.29, rRNA at 2.14, snoRNA at 2.12, tRNA at 1.97, and snRNA at 1.93. These values describe the magnitude of heterogeneity captured by SI in this analysis; they are not a ranking of biological complexity.

Across species, the profiles showed similar scale-dependent behavior, although absolute SI values varied among species and features. CDS and lncRNA illustrate the contrast: both changed across resolution, but their profiles occupied different regions of SI space.

Figure 1 and Figure 2 summarize the analytical framework and representative empirical observations, respectively.

**Figure 1:**
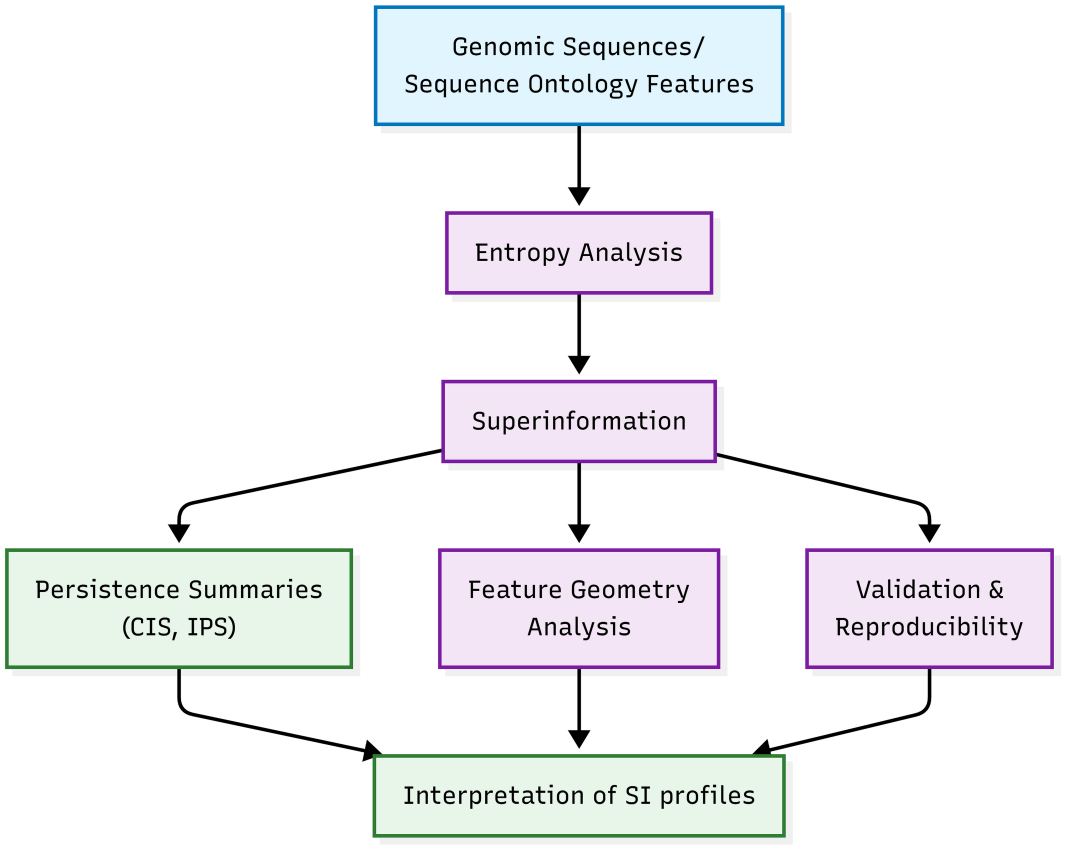
Study workflow and analytical framework. Reference genomic sequences and Sequence Ontology-defined genomic features were processed through block-level Shannon entropy and superinformation analysis. The resulting SI profiles were evaluated across sequence resolutions and characterized using complementary persistence summaries, feature-space geometry, validation and reproducibility analyses, and downstream interpretation of SI profiles.

**Figure 2:**
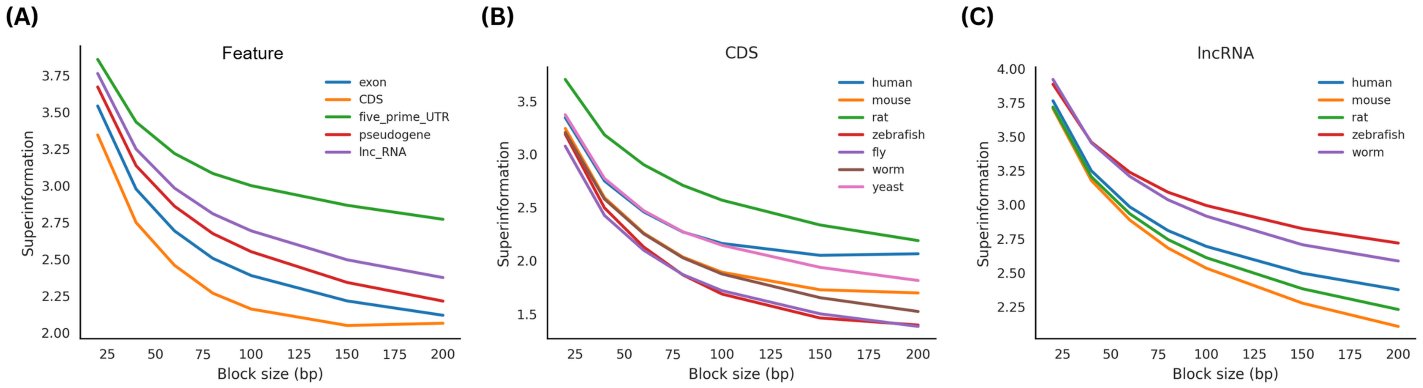
Representative multiscale superinformation profiles across genomic features and species. (A) Representative human superinformation profiles for exon, CDS, five-prime UTR, pseudogene, and lncRNA across block sizes from 20 to 200 bp. (B) Superinformation profiles for CDS across seven species: human, mouse, rat, zebrafish, fly, worm, and yeast. (C) Superinformation profiles for lncRNA across the species represented in the dataset: human, mouse, rat, zebrafish, and worm. The profiles illustrate feature-dependent differences in SI magnitude and changes in SI across sequence resolution.

### 3.2 Superinformation generally decreases with increasing block size while retaining feature-specific differences

Across block sizes from 20 to 200 bp, SI generally decreased as the sequence representation became coarser.

The decrease was common but not universal. Of the 86 evaluable feature-species trajectories, 72 were non-increasing across the seven block sizes (83.72%). The remaining trajectories contained one or more increases, so strict monotonicity cannot be assigned to every feature and species.

The feature classes also differed in both magnitude and persistence. lncRNA and untranslated regions, which had higher SI at 20 bp, generally retained higher SI at larger block sizes than CDS and several small RNA classes.

Normalizing to the 20-bp SI value made the feature profiles more similar. This suggests that part of the separation between features reflects overall SI magnitude rather than completely different responses to increasing block size.

Decay rates still differed. Coding sequences and several small RNA classes showed steeper early decreases, whereas lncRNA and untranslated regions declined more slowly. Thus, the profiles differed in both their initial SI magnitude and their rate of change with block size.

Within 20–200 bp, SI was generally highest at 20 bp, and no shared intermediate maximum was observed. The observed profiles therefore support a broad decline with resolution rather than a single common peak.

Figure 3 summarizes mean SI across feature classes and block sizes, normalized profiles for selected features, and the initial SI decay-rate distributions.

**Figure 3:**
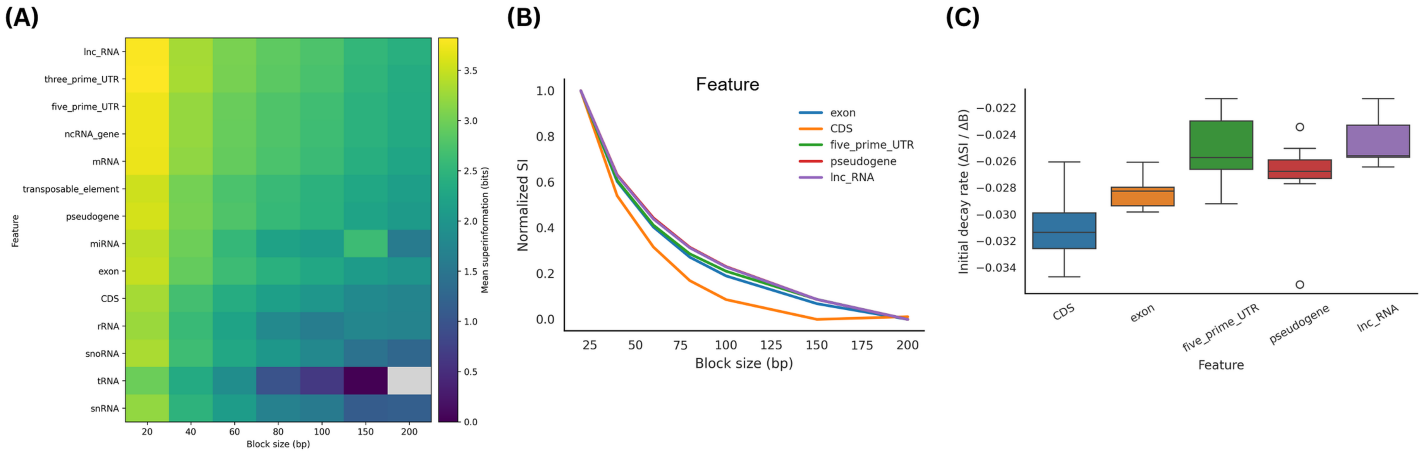
Resolution-dependent patterns in superinformation across genomic features. (A) Heatmap of mean SI (bits) across genomic feature classes and evaluated block sizes (20–200 bp). The gray cell indicates the unavailable tRNA value at 200 bp. (B) Normalized SI profiles for five selected feature classes (exon, CDS, five-prime UTR, pseudogene, and lncRNA), with each profile scaled to its SI value at 20 bp. (C) Initial SI decay rates for the same five feature classes, calculated from the change in SI between 20 and 40 bp as ΔSI*/*Δ*B*. More negative values indicate a steeper initial decline. Box plots show the distribution of available observations; points beyond the whiskers indicate outliers.

### 3.3 Observed superinformation generally exceeds the tested null expectations

We next compared the observed SI values with two simplified sequence-generation models: the block-count-controlled null and the species-level GC-controlled null.

Observed SI generally exceeded the block-count-controlled random null. The mean observed-minus-null difference was approximately 1.38 bits, the median was approximately 1.46 bits, and the largest difference was approximately 2.68 bits.

The excess was smaller under the species-level GC-controlled null but remained positive: the mean observed-minus-null difference was approximately 0.43 bits and the median was approximately 0.54 bits. Matching species-level GC composition therefore reduced the excess but did not reproduce the empirical SI profiles.

Associations with conventional sequence descriptors were modest. Mean SI correlated with mononucleotide Shannon entropy at *r* = −0.3224 (*R*^2^ = 0.1039), with dinucleotide entropy at *r* = −0.2574 (*R*^2^ = 0.0662), and with GC content at *r* = −0.3194 (*R*^2^ = 0.1020). No individual descriptor explained more than approximately 10.4% of the variation in mean SI.

The fast Monte Carlo procedure also closely matched the block-count-controlled null, with a correlation of approximately 0.999995 and differences of approximately 0.005 bits. Within the tested framework, the faster calculation therefore provides a close computational approximation.

Neither of the two tested random models reproduced the observed SI profiles, and no single baseline descriptor provided a strong explanation. The measured variation therefore contains structure not captured by these controls, but this does not establish independence from sequence composition or from other possible sequence-generating mechanisms.

Figure 4 shows the comparison with the block-count-controlled random null, the species-level GC-controlled random null, and the fast Monte Carlo approximation versus the block-count-controlled random null.

**Figure 4:**
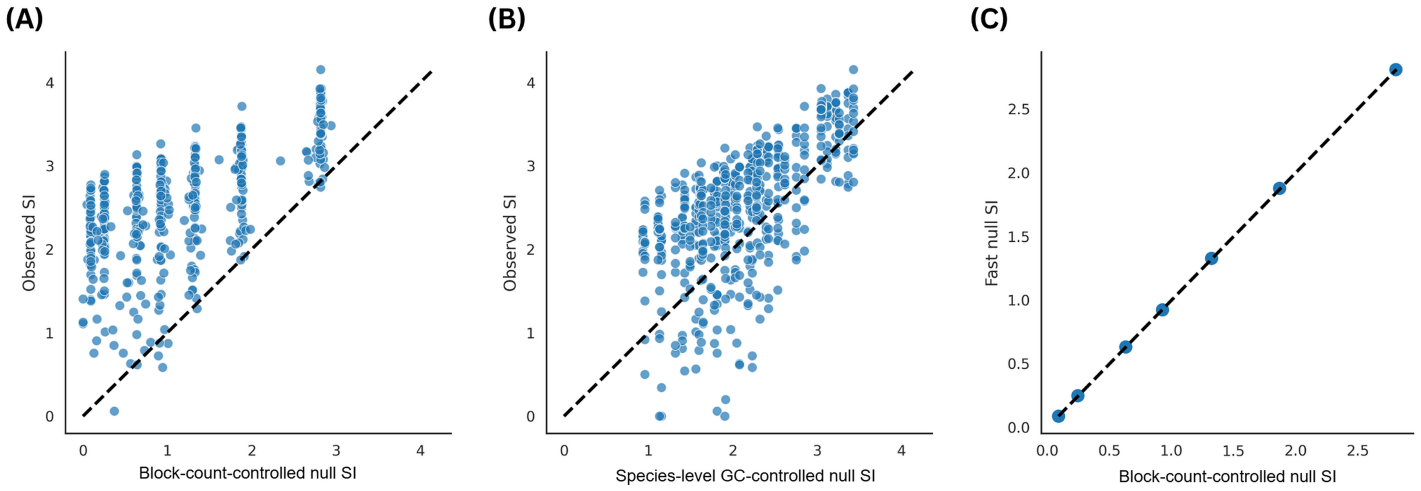
Comparison of observed superinformation with random null expectations. (A) Observed SI plotted against SI from the block-count-controlled random null, in which the number of simulated blocks was matched to the observed number of blocks for each species-feature-block-size combination. (B) Observed SI plotted against SI from the species-level GC-controlled random null, generated using the GC fraction of each species. In panels A and B, the dashed diagonal indicates equality between observed and null SI. (C) Comparison of the computationally efficient fast Monte Carlo null with the corresponding block-count-controlled random null estimates across the evaluated block sizes. The dashed diagonal indicates equality between the two estimates.

### 3.4 Multiscale SI profiles occupy a predominantly low-dimensional feature space

The retained SI profiles were represented as seven-dimensional vectors for geometric analysis.

PCA of the unscaled 13 ×7 matrix concentrated most of the variance in the first two components. PC1 explained 93.341% and PC2 explained 4.518%, for 97.859% combined variance.

The dominance of PC1 indicates that much of the variation is coordinated across block sizes: profiles with consistently higher SI tend to separate from profiles with consistently lower SI along this axis.

UMAP provided a complementary nonlinear view of the same 13 complete profiles. The canonical two-dimensional embedding was regenerated from the frozen feature-level SI matrix using Euclidean distance, 10 neighbors, a minimum distance of 0.2, and random state 42. The resulting spatial structure describes similarity in multiscale SI behavior and was not treated as a biological classification.

The same distinction applies to the broader feature groups shown in the visualization: their positions reflect the SI profiles, while biological interpretation requires independent genomic annotation and other sequence properties.

Figure 5 presents the UMAP and PCA representations of the SI profile space.

**Figure 5:**
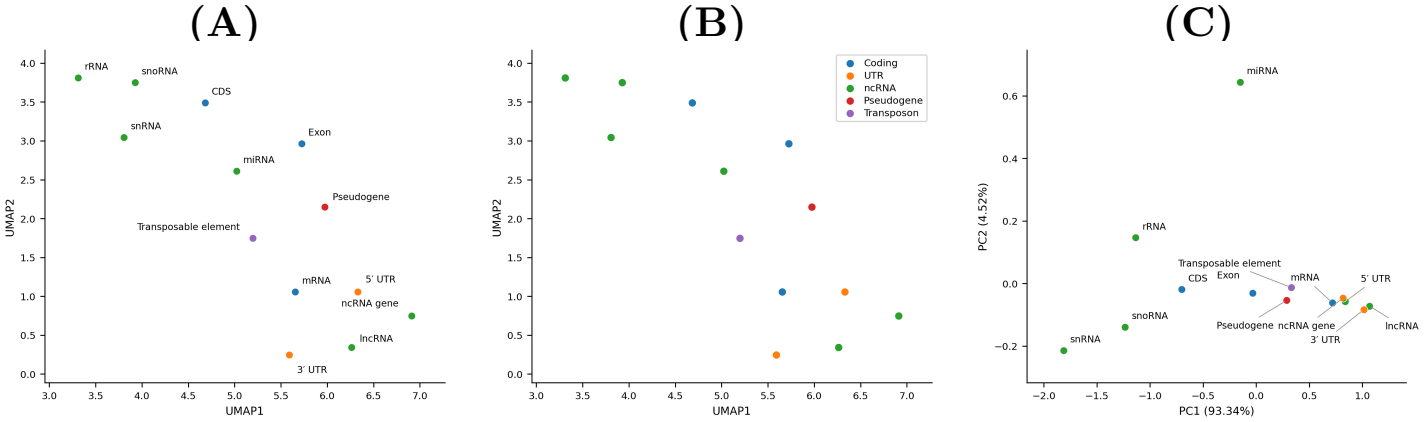
Low-dimensional representation of multiscale superinformation profiles. The study included 14 feature types overall; 13 complete feature-level profiles were retained for geometry after averaging SI across available species at each of seven block sizes, without imputation or scaling. (A) Feature-labelled UMAP. (B) The same canonical UMAP coordinates colored by the descriptive visualization groups Coding, UTR, ncRNA, Pseudogene, and Transposon; these groups are not inferred classes. UMAP was regenerated from the frozen feature-level matrix in two dimensions using Euclidean distance, `n neighbors=10`, `min dist=0.2`, and `random state=42`. (C) PCA of the same unscaled 13 7× matrix. PC1 and PC2 explain 93.341% and 4.518% of the variance, respectively, for a cumulative 97.859%.

### 3.5 Hierarchical organization of feature profiles

Hierarchical clustering was applied to the same 13 complete feature-level mean SI profiles using average linkage and Euclidean distance on the unscaled 13 ×7 matrix. Four clusters were obtained with the `maxclust` criterion.

The dendrogram grouped features with similar multiscale SI trajectories. Cluster 1 contained CDS and rRNA; Cluster 2 contained snRNA and snoRNA; Cluster 3 contained exon, five-prime UTR, lncRNA, mRNA, ncRNA gene, pseudogene, three-prime UTR, and transposable element; and Cluster 4 contained miRNA. The assignments describe quantitative similarity among SI profiles, not biological or ontology classes.

Figure 6 shows the hierarchical organization of the feature profiles and their saved cluster memberships.

**Figure 6:**
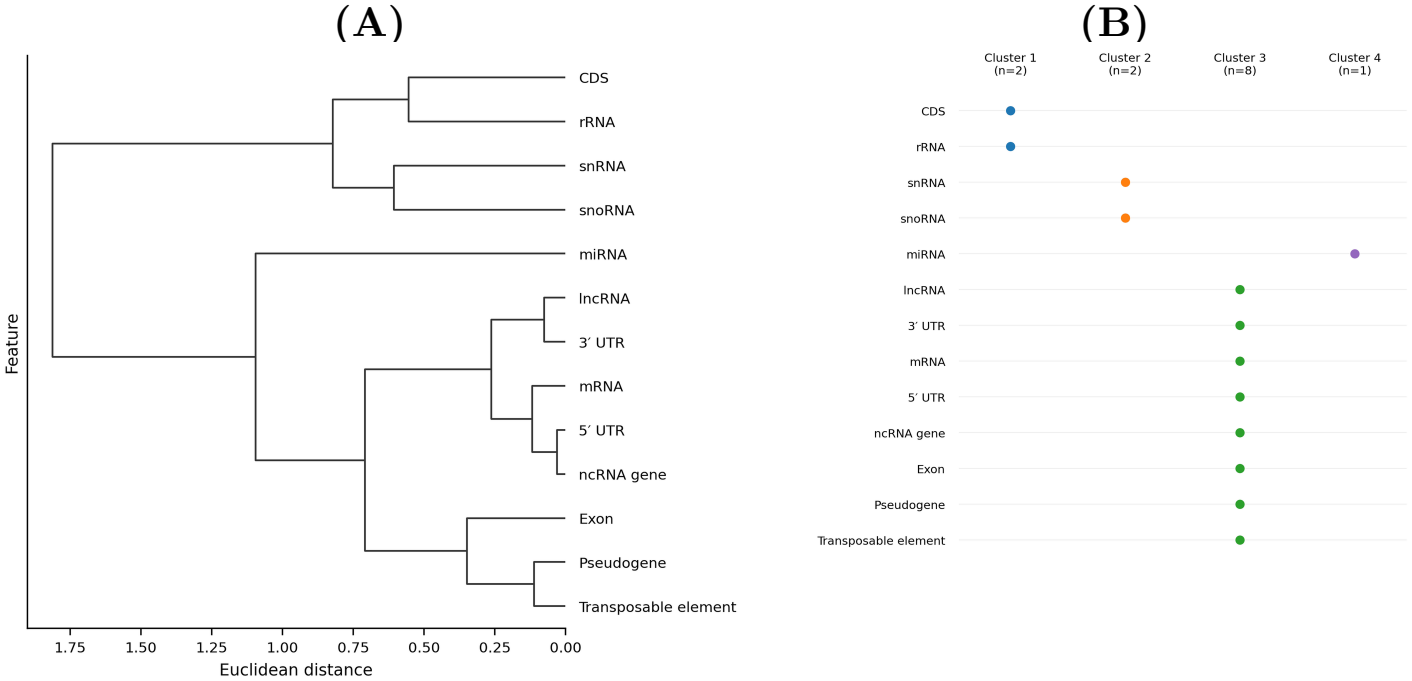
Hierarchical organization of genomic features based on multiscale superinformation profiles. (A) Horizontal dendrogram from average-linkage hierarchical clustering of the unscaled 13× 7 complete feature-level SI matrix using Euclidean distance. Each feature profile was obtained by averaging SI across available species at the seven evaluated block sizes. (B) Feature-by-cluster membership matrix for the four saved `maxclust` assignments. Each feature belongs to exactly one cluster. The clustering describes quantitative similarity in multiscale SI trajectories and does not represent a biological or functional classification.

### 3.6 Characteristic Information Scale and information persistence

Two complementary measures were used to summarize the multiscale profiles: a threshold-crossing scale and cumulative persistence across the evaluated resolution range.

The Characteristic Information Scale (CIS) was defined as the block size at which SI falls to 50% of its 20-bp value. Finite CIS estimates were obtained for seven feature classes: 181.98 bp for miRNA (*n* = 1), 148.92 bp for pseudogene (*n* = 1), 143.49±24.41 bp for CDS (*n* = 3), 92.31±48.59 bp for snRNA (*n* = 6), 85.62 ± 12.42 bp for snoRNA (*n* = 4), 72.93 ± 19.18 bp for rRNA (*n* = 4), and 65.26± 3.10 bp for tRNA (*n* = 3). These values are descriptive and depend on feature availability across species.

For exon, five-prime UTR, lncRNA, mRNA, ncRNA gene, three-prime UTR, and transposable element, the 50% threshold was not reached within the 20– 200 bp window. These features therefore have no finite CIS estimate within the study window; this does not locate a characteristic scale beyond 200 bp.

The Information Persistence Score (IPS) was calculated by trapezoidal integration over the available evaluated block sizes within the 20–200 bp study window. Complete trajectories span the full window; incomplete trajectories contribute only over their available evaluated points. Mean normalized IPS was highest for lncRNA (133.07 ±5.95), followed by five-prime UTR (131.34 ±8.44), ncRNA gene (131.29 ±5.12), three-prime UTR (131.01± 6.54), and mRNA (129.74± 7.04). Transposable element (128.72 ±10.94), pseudogene (125.52 ±8.44), and exon (122.95 ±3.21) also had relatively high values. Lower values were observed for CDS (114.81± 9.05), rRNA (106.04 ±23.99), snoRNA (100.26 ±21.43), snRNA (87.39 ±31.42), miRNA (66.19± 55.62), and tRNA (39.03± 7.46).

The IPS values broadly followed the differences seen in the SI profiles. Features with higher SI across multiple block sizes generally accumulated larger integrated scores, whereas lower or more rapidly attenuated profiles produced smaller values. CIS records a specified threshold crossing, while IPS summarizes cumulative SI over the available evaluated span.

CIS and IPS therefore capture complementary aspects of resolution-dependent behavior: CIS describes a threshold scale within the tested range, whereas IPS summarizes cumulative SI across the available range.

Additional supporting analyses are provided in the Supplementary Information, including the complete species–feature SI trajectories (Supplementary Fig. S1), PCA of the feature-level profiles (Supplementary Fig. S2), null-model convergence (Supplementary Fig. S3), entropy-variance analyses (Supplementary Figs. S4–S5), the species-level GC-controlled null comparison (Supplementary Fig. S6), baseline sequence-descriptor associations (Supplementary Fig. S7), and feature availability and CIS/IPS summaries (Supplementary Tables S1–S2).

## 4 Discussion

### 4.1 Extending the superinformation framework across genomic features and scales

This study extends the superinformation framework introduced by Bose and Chouhan (2011) without changing its underlying definition. Their measure takes the entropy of the distribution of local Shannon entropy values from sequence blocks, capturing heterogeneity among local information states rather than only average nucleotide composition. They showed that it could distinguish coding and noncoding DNA and could also be applied to untranslated regions. We apply the same framework across 14 Sequence Ontology-defined feature classes, seven species, and seven block sizes from 20 to 200 bp.

Here SI is treated as a profile across resolution rather than as a value at one selected block size. This is relevant because sequence analyses can change with scale: local Shannon entropy varies along DNA, compositional segmentation identifies heterogeneous domains, and comparative sequence statistics can depend on representation scale or word length (Bussi et al., 2021; Cristadoro et al., 2018; Moya et al., 2020; Thanos et al., 2018). Tracking SI as block size changes therefore places the measure in that multiscale setting.

The profiles differed among genomic feature classes, both in SI magnitude and in their trajectories across the tested resolutions. The contribution here is therefore not a new definition of superinformation, but a broader characterization of what the existing measure shows when applied systematically across genomic categories and scales.

### 4.2 Feature-dependent superinformation reflects differences in sequence heterogeneity

The separation among feature classes means that the distribution of local sequence entropy differs among the genomic categories examined here. lncRNA and untranslated regions occupied higher-SI regimes across much of the range, while CDS and several compact RNA classes showed lower SI and faster attenuation with increasing block size. Because SI is the entropy of the distribution of block-level entropies, the most direct interpretation is a difference in the heterogeneity of local sequence variability.

These observations are consistent with earlier work showing that genomic sequence composition is not spatially uniform. Local Shannon entropy can identify spatial fluctuations in sequence complexity, and compositional segmentation can distinguish domains with different local properties despite similar genome-wide averages (Moya et al., 2020; Thanos et al., 2018). Superstatistical analyses have also shown that similar global GC content can coexist with correlated local compositional states (Bogachev et al., 2017). These findings provide context for the different SI profiles observed here, but do not establish their biological cause.

SI alone does not determine why these differences occur. A higher SI value should not be read as greater biological complexity, functional importance, or evolutionary constraint. The feature ordering here reflects the magnitude of local entropy heterogeneity captured under the specified analysis. This is especially relevant for classes such as pseudogenes and transposable elements, which can contain mixtures of repetitive, inherited, divergent, and functionally co-opted sequence.

Normalization by the 20-bp value reduced much of the separation among feature classes, so part of the difference reflects overall SI magnitude. Initial decay rates still differed, showing that the classes also differ in how their measured heterogeneity changes with block size. The multiscale profile therefore contains information beyond the initial SI value.

### 4.3 Resolution dependence is an intrinsic component of the measurement

SI generally declined as block size increased: 72 of 86 evaluable trajectories were non-increasing across the seven tested resolutions. The remaining trajectories show that the pattern is not universal. Even so, the predominance of decreasing profiles indicates that the distribution of local entropy values usually changes as the sequence is represented with progressively larger blocks.

This pattern is consistent with broader sequence analyses in which genomic organization depends on the scale of observation. Local entropy, segmentation, and comparative *k*-mer analyses can expose different aspects of sequence organization (Bussi et al., 2021; Moya et al., 2020; Thanos et al., 2018). Cristadoro et al. (2018) also reported multiple characteristic regimes in human genomic sequence statistics, providing another example of scale-dependent organization. These observations are limited to the 20–200 bp window. They do not show whether similar behavior continues at kilobase, megabase, or chromosome scales.

Likewise, the absence of a shared intermediate SI maximum in this range does not rule out longer-range maxima or transitions outside the study window.

Block size is therefore an analytical resolution parameter rather than only an implementation choice. A single block size gives one view of the SI land-scape, whereas the full profile records how measured heterogeneity changes under coarse-graining.

### 4.4 The null models constrain, but do not exhaust, explanations for SI

The null models provide a control on the interpretation of the feature-dependent SI differences. Observed SI exceeded the block-count-controlled null by approximately 1.38 bits on average and remained elevated relative to the species-level GC-controlled null by approximately 0.43 bits on average. Thus, the observed profiles were not reproduced by the specific random sequence models tested here.

The smaller excess under the GC-controlled null shows that nucleotide composition contributes to the signal. The remaining difference means that species-level GC composition alone does not reproduce the empirical distribution of block-level entropy values. This is consistent with evidence that global composition can coexist with substantial spatial variation in local sequence properties (Bogachev et al., 2017; Moya et al., 2020) and that local sequence variation can contain information beyond mean GC content or whole-region summaries (López-Cortegano et al., 2021).

The baseline analyses support the same limited conclusion. Mean SI was only modestly associated with mononucleotide entropy, dinucleotide entropy, and GC content; the largest individual *R*^2^ was approximately 0.104. This does not show that SI is independent of sequence composition. Rather, none of the individual descriptors tested provides a sufficient one-dimensional explanation of the observed SI variation.

The null results are therefore evidence against the specific simplified alternatives tested, not against all possible sequence-generating mechanisms. More detailed null models could include feature-specific composition, higher-order dependencies, repeat structure, or spatial correlations and would provide stronger tests of what contributes to the SI landscape.

### 4.5 Multiscale SI profiles define a structured quantitative feature space

The 13 complete seven-dimensional SI profiles also showed coordinated structure. PC1 and PC2 captured 97.859% of the variance, with PC1 accounting for 93.341% and PC2 for 4.518%. This concentration is consistent with many feature profiles differing mainly in overall SI magnitude while retaining broadly similar trajectories across resolution.

UMAP gave a complementary view of the same profiles, which occupied structured regions of the embedding, and hierarchical clustering grouped features with similar multiscale SI trajectories. Together, these analyses organize the profiles by quantitative similarity in their behavior across resolution.

The geometric results are descriptive. Similarity in SI space does not establish shared biological function, common evolutionary history, or membership in an externally defined biological class. The analysis describes a quantitative organization of sequence-heterogeneity profiles, not a biological classification.

This interpretation is consistent with sequence-complexity and entropy-based work showing that genomes contain structured distributions of local sequence states rather than being fully described by single global summaries (Bogachev et al., 2017; Moya et al., 2020; Thanos et al., 2018). Here, the object of comparison is the resolution-dependent SI profile itself.

### 4.6 Scope, limitations, and future directions

Several limitations set the scope of these conclusions. The analysis used seven reference genomes and 14 selected Sequence Ontology feature classes, and feature availability differed among species and categories. Some SI trajectories also did not extend to every resolution. The results therefore describe the curated panel and the observations available within these genomes, not all genomic sequence classes.

The block sizes ranged from 20 to 200 bp. This range addresses fine-scale sequence organization but not larger compositional or regulatory domains. Extending the analysis to broader spatial scales would test whether additional transitions or characteristic regimes occur beyond this range.

The null models were intentionally simplified. The block-count-controlled and species-level GC-controlled models test specific alternatives but do not reproduce the full dependence structure of genomic sequence. The conclusion is therefore relative: the observed profiles contain structure not reproduced by these controls.

SI also summarizes the distribution of local entropy values and therefore discards information by construction. Sequences with different nucleotide identities, positional arrangements, or motif grammars can produce similar local-entropy distributions. SI should therefore complement, rather than replace, sequence representations that retain higher-order word identity or positional relationships.

The geometric analyses are likewise descriptive and use a relatively small number of feature-level profiles. PCA, UMAP, and hierarchical clustering describe profile similarity but do not provide independent biological validation. The results should therefore be kept separate from biological annotation.

Future work can extend the comparison in several directions. Larger comparative datasets could test whether SI profiles recur across additional taxa and assemblies. A wider block-size range could connect fine-scale SI behavior to broader compositional organization. More structured null models could test higher-order nucleotide dependencies and spatial correlations. Combining SI with independent sequence, epigenomic, or functional measurements could also help determine which components of the SI landscape correspond to specific biological processes, rather than assigning biological meaning from SI alone.

## Author Contributions

Sarvesh Galgale: Data curation, formal analysis, investigation, methodology, software, validation, visualization, writing (original draft, review and editing). Praharshit Sharma: Conceptualization.

## Data and Code Availability

The data, analysis scripts, and supporting computational materials associated with this study are publicly available at https://github.com/sarvezh1/superinformation-genomics. The repository has been archived on Zenodo under DOI https://doi.org/10.5281/zenodo.24437905 (Version 1.0.0). The repository includes the computational workflows and supporting materials associated with the analyses presented in this study.

## Funding

This research received no external funding.

## Conflict of Interest

The authors declare no competing interests.

## Ethics Statement

Ethics approval was not applicable to this study, as it used publicly available reference genome sequences and annotations and did not involve new experiments on human participants or animals.

## Notes

### Competing Interest Statement

The authors have declared no competing interest.

