## Supplementary Information for "Multiscale superinformation reveals structured organization of genomic sequence features"

### Supplementary Methods

#### S1. Additional multiscale superinformation profiles

Superinformation (SI) was evaluated across seven block sizes of 20, 40, 60, 80, 100, 150, and 200 bp using the procedure described in the main manuscript. The complete set of species–feature trajectories is shown here to provide the full multiscale profile dataset underlying the representative profiles presented in the main manuscript. Each trajectory corresponds to one species–feature combination represented in the final dataset.

#### S2. Feature-space analysis

The study included 14 genomic feature types overall. For geometric analyses, SI values were averaged across the available species for each genomic feature at each block size. Thirteen feature types had complete values at all seven resolutions and were retained; the incomplete tRNA profile was excluded. No missing values were imputed or interpolated. Principal component analysis was performed directly on the resulting unscaled 13 × 7 feature-by-block-size matrix, preserving differences in SI magnitude across resolutions. PC1 explained 93.341% of the variance and PC2 explained 4.518%, for a cumulative 97.859%. The same matrix was used for the canonical two-dimensional UMAP visualization with Euclidean distance, `n_neighbors=10`, `min_dist=0.2`, and `random_state=42`.

#### S3. Null-model convergence

The block-count-controlled random null generated independent random DNA blocks in a number equal to the observed number of blocks for each species–feature–block-size combination, using equal probabilities for A, C, G, and T. The fast Monte Carlo null used 50,000 random blocks at each evaluated block size with uniform nucleotide probabilities. The two approaches produced nearly identical SI estimates, with a correlation of approximately 0.999995 and differences of approximately 0.005 bits.

#### S4. Entropy variance analyses

Additional analyses examined the variability of block-level Shannon entropy across sequence resolution. These analyses provide complementary characterization of how local entropy variance changes with block size and how entropy variance relates to superinformation. Entropy variance was calculated as the population variance across contributing block-entropy values, using division by the number of blocks. For each genomic feature and block size, mean variance was calculated across species with available observations. Shaded bands in Supplementary Fig. S4 represent 95% bootstrap confidence intervals around these means, based on 1,000 bootstrap resamples.

#### S5. Species-level GC-controlled random null analysis

A complementary species-level GC-controlled random null was used to assess whether observed SI could be reproduced using species-specific nucleotide composition. For each species and block size, 50,000

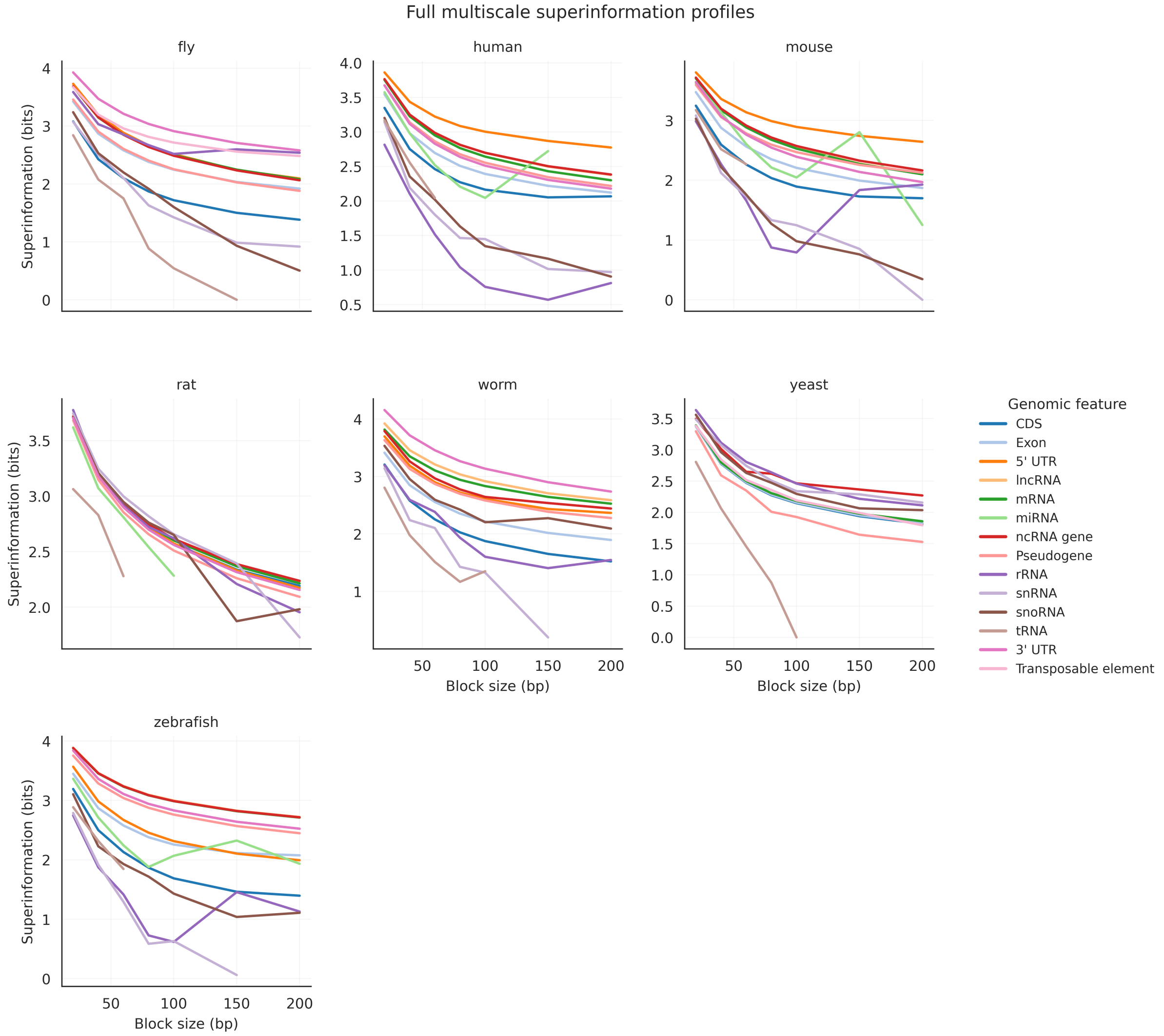


**Supplementary Figure S1. Full multiscale superinformation profiles across species and genomic feature classes.** Superinformation values are shown across the seven evaluated block sizes for the available genomic feature classes in human, mouse, rat, zebrafish, fly, worm, and yeast. Each coloured trajectory represents one Sequence Ontology feature class. The same feature-to-colour mapping is used across all species panels; trajectories are shown where data are available.

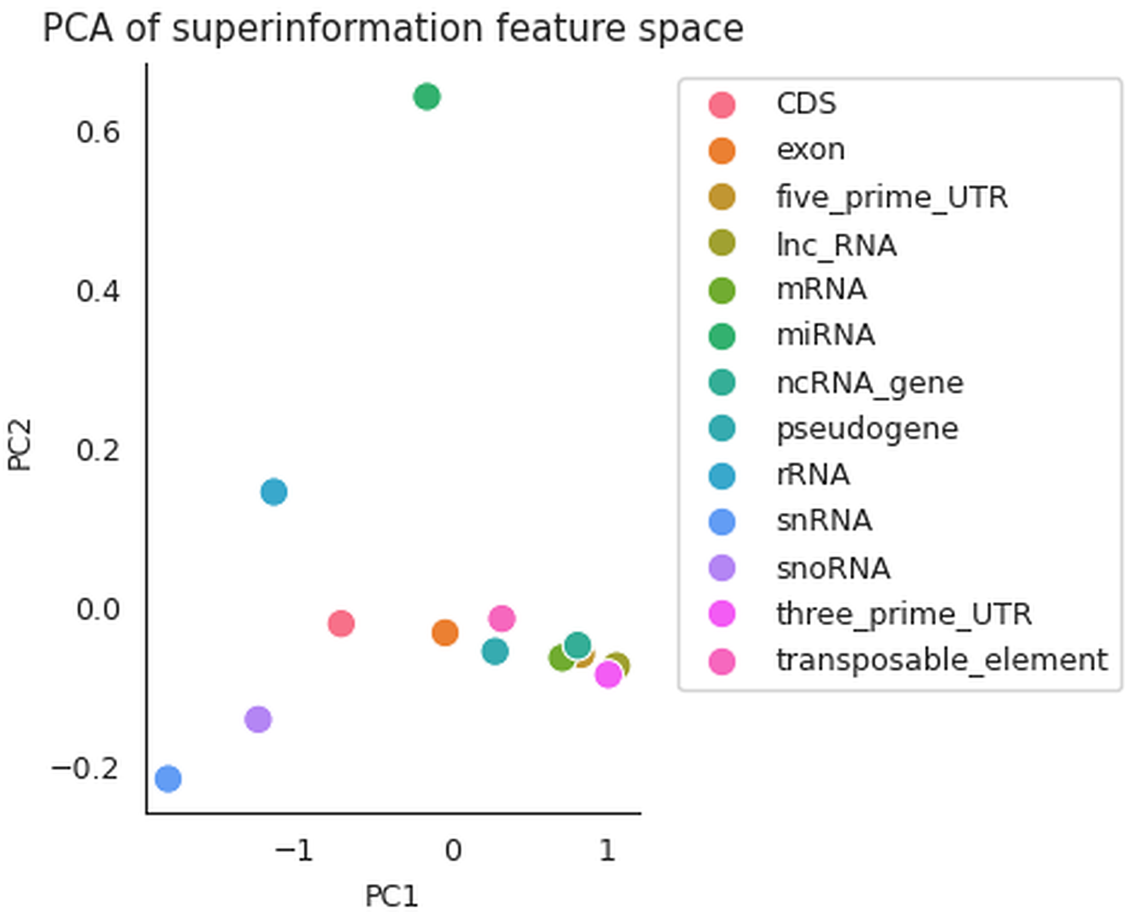


**Supplementary Figure S2. Principal component representation of multiscale superinformation profiles.** Each point represents one of the 13 complete genomic feature profiles retained from the 14-feature study after averaging SI across available species at each of the seven evaluated block sizes. PCA was performed directly on the unscaled $13 \times 7$ feature-by-block-size matrix. PC1 and PC2 explain 93.341% and 4.518% of the variance, respectively, for a cumulative 97.859%.

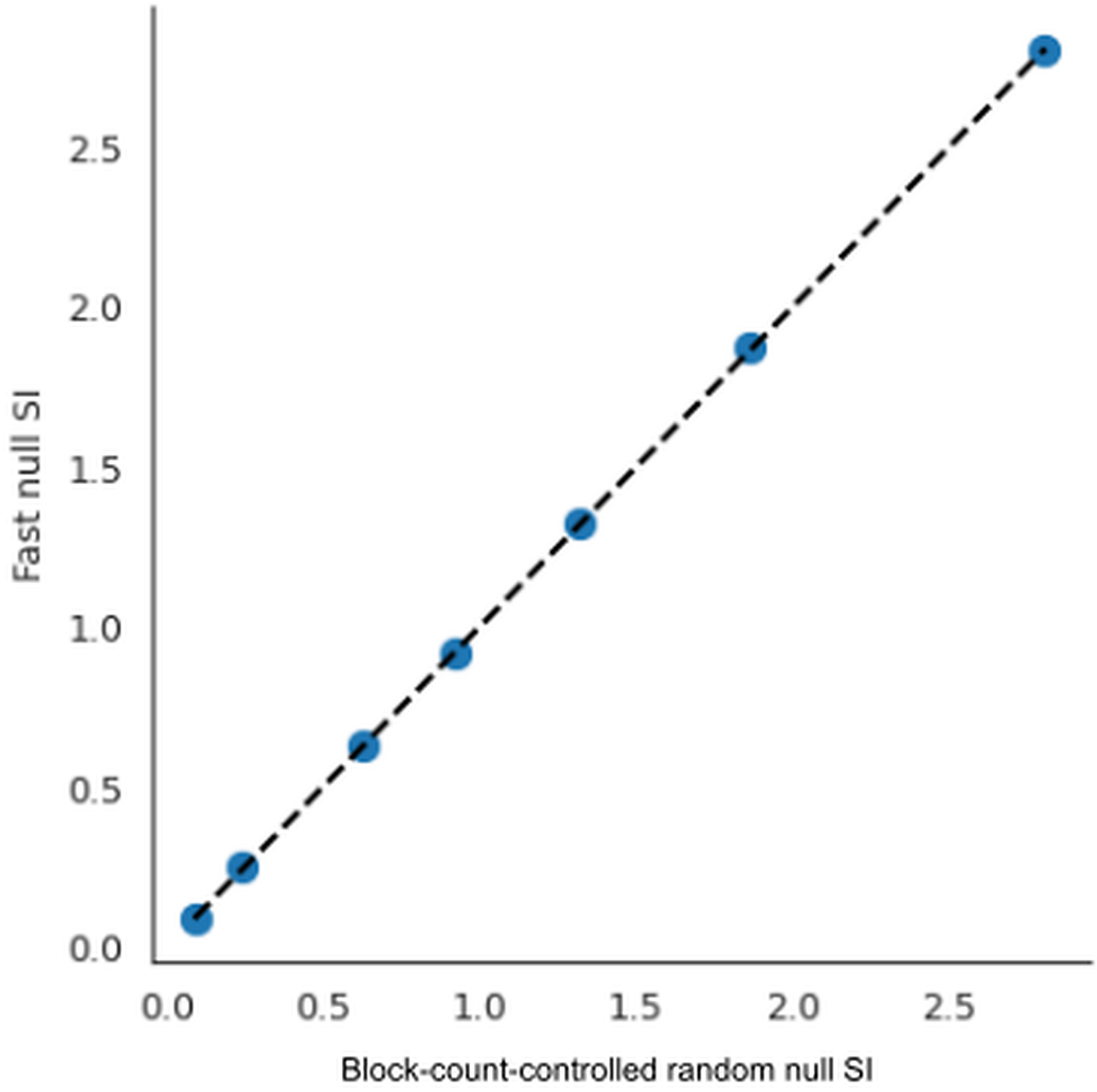


**Supplementary Figure S3. Agreement between fast and block-count-controlled Monte Carlo null estimates.** Superinformation estimated using the fast Monte Carlo procedure at each of the seven block sizes is compared with the corresponding block-size-aggregated block-count-controlled random-null estimate. The dashed diagonal indicates equality between the two estimates.

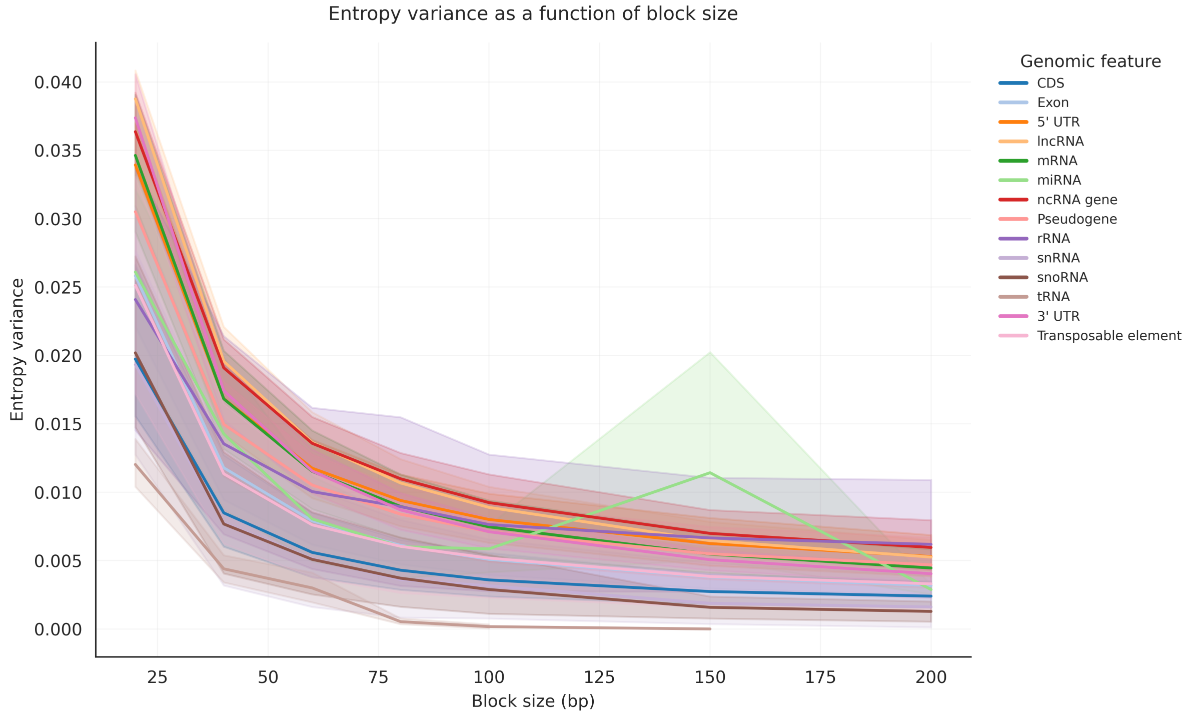


**Supplementary Figure S4. Entropy variance as a function of block size.** Each coloured line represents the mean variance of block-level Shannon entropy for one genomic feature class across the species with available observations at each block size. Shaded bands indicate the 95% bootstrap confidence intervals around the corresponding means.

independent random DNA blocks were generated using the species-level GC fraction to define the nucleotide probabilities, with A and T assigned equal probability and C and G assigned equal probability. Superinformation was then calculated from the resulting distribution of block-level entropy values.

The GC-controlled null reduced the observed-minus-null SI excess relative to the block-count-controlled null but did not reproduce the empirical SI profiles. The mean observed-minus-null difference was approximately 0.43 bits, with a median difference of approximately 0.54 bits.

#### S6. Baseline sequence-descriptor analysis

To assess relationships between superinformation and conventional sequence descriptors, feature-level mean SI was compared with mononucleotide Shannon entropy, dinucleotide entropy, and GC content. Feature-level SI was summarized from the available rows of the 578-observation SI dataset. The baseline descriptor dataset was summarized separately from 89 species–feature rows and used up to 1,000 sequences per species–feature row.

Pearson correlation coefficients were calculated between mean SI and each descriptor. The resulting associations were modest: mononucleotide entropy, $r = -0.3224$ ($R^2 = 0.1039$); dinucleotide entropy, $r = -0.2574$ ($R^2 = 0.0662$); and GC content, $r = -0.3194$ ($R^2 = 0.1020$). No individual descriptor explained more than approximately 10.4% of the variation in mean SI.

#### S7. Data availability and reproducibility

The final SI dataset contains 578 species–feature–block-size observations covering seven species, 14 genomic feature classes, and seven evaluated block sizes. The supplementary analyses use the same final SI values and derived analysis outputs used for the main manuscript.

The data, analysis scripts, and supporting computational materials are publicly available at `https://github.com/sarvezh1/superinformation-genomics`. The archived release (Version 1.0.0) is available on Zenodo at `https://doi.org/10.5281/zenodo.24437905`.

The computational workflow was implemented in Python using standard scientific-computing libraries. The analysis framework was packaged as a reusable software module and validated against

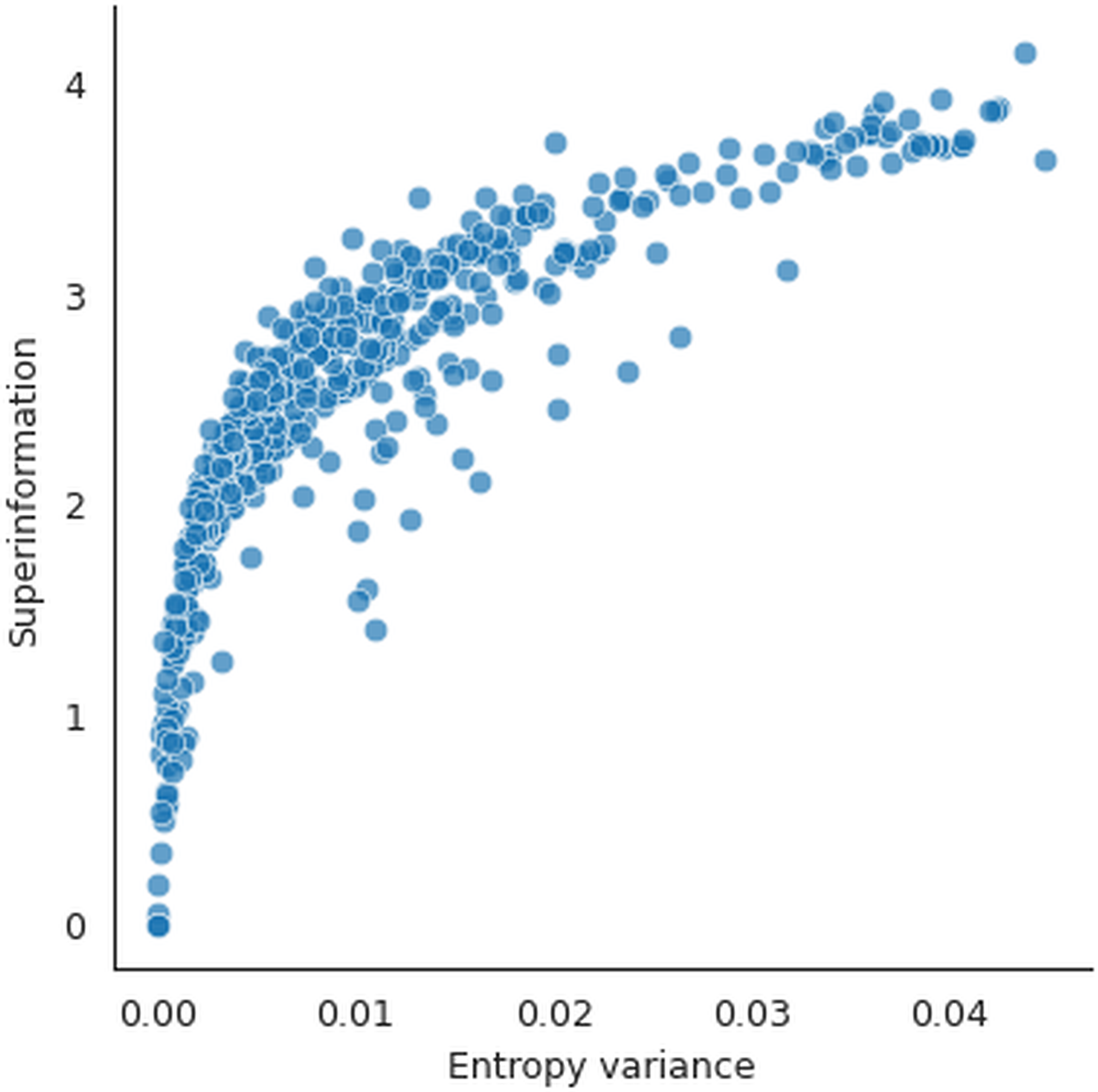


**Supplementary Figure S5. Relationship between entropy variance and superinformation.** Block-level Shannon entropy variance is compared with superinformation across the evaluated observations. The analysis provides a complementary view of the variability in local entropy underlying the SI statistic.

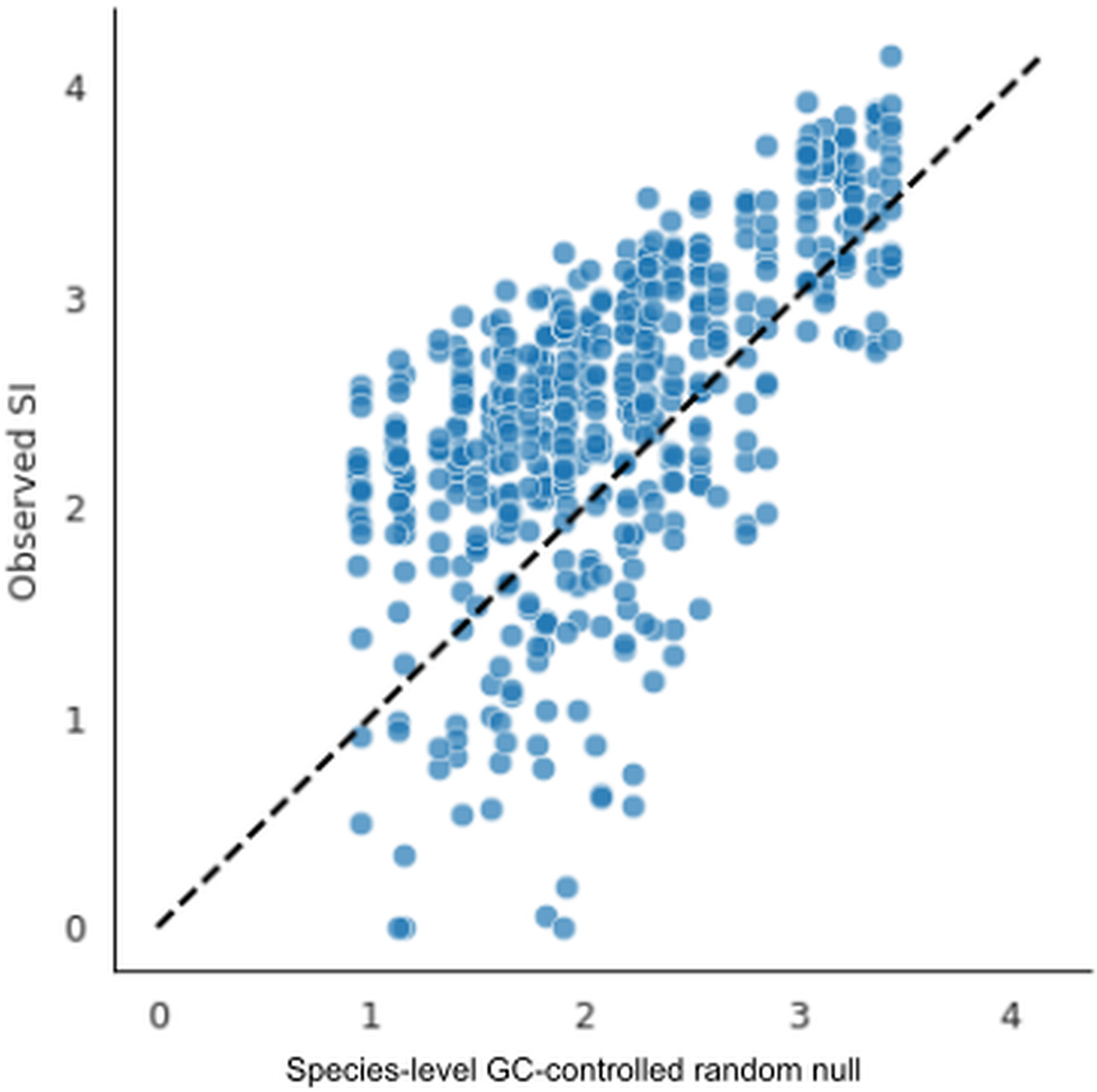


**Supplementary Figure S6. Comparison with the species-level GC-controlled random null.** Observed superinformation is plotted against superinformation estimated from the species-level GC-controlled random null. The dashed diagonal indicates equality between observed and null SI. The null preserves the species-level GC fraction while generating independent random DNA blocks.

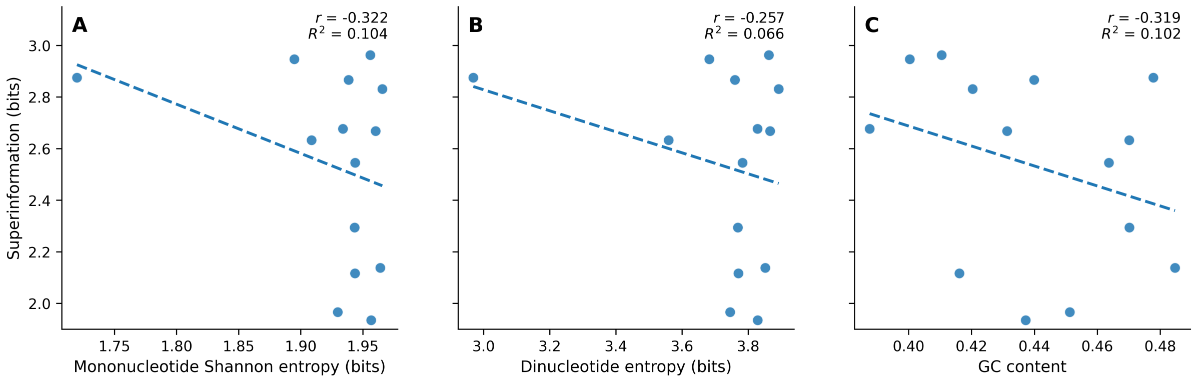


**Supplementary Figure S7. Relationships between superinformation and conventional sequence descriptors.** (A) Feature-level superinformation plotted against mononucleotide Shannon entropy. (B) Superinformation plotted against dinucleotide entropy. (C) Superinformation plotted against GC content. Dashed lines represent linear regression fits. Pearson correlation coefficients and corresponding coefficients of determination are shown in each panel.

retained reference outputs. Reproduction of the CIS calculations using the validated implementation recovered the reference dataset structure and finite-value coverage; across the 22 finite CIS values, the maximum absolute difference was $1.42 \times 10^{-14}$.

The feature-by-resolution SI outputs, baseline descriptor data, null-model outputs, CIS values, and IPS values are retained as analysis outputs.

### Supplementary Tables

**Supplementary Table S1. Genomic feature classes, broad feature groups, and available sequence resolutions across species.**

Entries indicate the number of evaluated block sizes available for each species–feature combination, out of seven possible resolutions (20, 40, 60, 80, 100, 150, and 200 bp). Zero indicates that the species–feature combination was not represented in the final SI dataset. Broad feature groups are used only for visualization and do not represent biological ontology categories.

| Feature | Broad class | Human | Mouse | Rat | Zebrafish | Fly | Worm | Yeast |
|---|---|---|---|---|---|---|---|---|
| CDS | Coding | 7 | 7 | 7 | 7 | 7 | 7 | 7 |
| exon | Coding | 7 | 7 | 7 | 7 | 7 | 7 | 7 |
| five_prime_UTR | UTR | 7 | 7 | 7 | 7 | 7 | 7 | 0 |
| lnc_RNA | ncRNA | 7 | 7 | 7 | 7 | 0 | 7 | 0 |
| mRNA | Coding | 7 | 7 | 7 | 7 | 7 | 7 | 7 |
| miRNA | ncRNA | 6 | 7 | 5 | 7 | 1 | 1 | 0 |
| ncRNA_gene | ncRNA | 7 | 7 | 7 | 7 | 7 | 7 | 7 |
| pseudogene | Pseudogene | 7 | 7 | 7 | 7 | 7 | 7 | 7 |
| rRNA | ncRNA | 7 | 7 | 7 | 7 | 7 | 7 | 7 |
| snRNA | ncRNA | 7 | 7 | 7 | 6 | 7 | 6 | 7 |
| snoRNA | ncRNA | 7 | 7 | 7 | 7 | 7 | 7 | 7 |
| tRNA | ncRNA | 3 | 3 | 3 | 3 | 6 | 5 | 5 |
| three_prime_UTR | UTR | 7 | 7 | 7 | 7 | 7 | 7 | 0 |
| transposable_element | Transposon | 0 | 0 | 0 | 0 | 7 | 0 | 7 |

**Supplementary Table S2. Characteristic Information Scale and Information Persistence Score across genomic feature classes.**

CIS denotes the Characteristic Information Scale, defined as the block size at which SI reaches 50% of its value at 20 bp. CIS values are reported as mean ± SD across available finite species–feature observations; "NR" indicates that the 50% threshold was not reached within the 20–200 bp evaluation window. Raw IPS denotes the trapezoidal integral of SI over the available evaluated block sizes within the 20–200 bp study window and has units of bit·bp. Complete trajectories span 20–200 bp; incomplete trajectories are integrated only over their available evaluated block sizes within that window. Normalized IPS applies the same available-point integration after normalization by SI(20) and therefore has units of bp. $n_{\text{CIS}}$ and $n_{\text{IPS}}$ indicate the number of finite observations contributing to each summary.

| Feature | $n_{\text{CIS}}$ | CIS (bp) | $n_{\text{IPS}}$ | Raw IPS (bit·bp) | Normalized IPS (bp) |
|---|---|---|---|---|---|
| lnc_RNA | 0 | NR | 5 | 506.18 ± 34.93 | 133.07 ± 5.95 |
| three_prime_UTR | 0 | NR | 6 | 501.93 ± 49.42 | 131.01 ± 6.54 |
| five_prime_UTR | 0 | NR | 6 | 490.28 ± 43.81 | 131.34 ± 8.44 |
| ncRNA_gene | 0 | NR | 7 | 488.89 ± 30.26 | 131.29 ± 5.12 |
| mRNA | 0 | NR | 7 | 481.51 ± 44.73 | 129.74 ± 7.04 |
| transposable_element | 0 | NR | 2 | 455.23 ± 65.01 | 128.72 ± 10.94 |
| pseudogene | 1 | 148.92 | 7 | 451.00 ± 48.03 | 125.52 ± 8.44 |
| exon | 0 | NR | 7 | 428.61 ± 24.08 | 122.95 ± 3.21 |
| CDS | 3 | 143.49 ± 24.41 | 7 | 381.17 ± 52.56 | 114.81 ± 9.05 |
| rRNA | 4 | 72.93 ± 19.18 | 7 | 352.45 ± 119.18 | 106.04 ± 23.99 |
| snoRNA | 4 | 85.62 ± 12.42 | 7 | 339.39 ± 97.20 | 100.26 ± 21.43 |
| snRNA | 6 | 92.31 ± 48.59 | 7 | 288.05 ± 129.41 | 87.39 ± 31.42 |
| miRNA | 1 | 181.98 | 6 | 233.74 ± 195.15 | 66.19 ± 55.62 |
| tRNA | 3 | 65.26 ± 3.10 | 7 | 114.86 ± 17.47 | 39.03 ± 7.46 |
